# Genomic prediction in the wild: A case study in Soay sheep

**DOI:** 10.1101/2020.07.15.205385

**Authors:** B Ashraf, DC Hunter, C Bérénos, PA Ellis, SE Johnston, JG Pilkington, JM Pemberton, J Slate

## Abstract

Genomic prediction, the technique whereby an individual’s genetic component of their phenotype is estimated from its genome, has revolutionised animal and plant breeding and medical genetics. However, despite being first introduced nearly two decades ago, it has hardly been adopted by the evolutionary genetics community studying wild organisms. Here, genomic prediction is performed on eight traits in a wild population of Soay sheep. The population has been the focus of a >30 year evolutionary ecology study and there is already considerable understanding of the genetic architecture of the focal Mendelian and quantitative traits. We show that the accuracy of genomic prediction is high for all traits, but especially those with loci of large effect segregating. Five different methods are compared, and the two methods that can accommodate zero-effect and large-effect loci in the same model tend to perform best. If the accuracy of genomic prediction is similar in other wild populations, then there is a real opportunity for pedigree-free molecular quantitative genetics research to be enabled in many more wild populations; currently the literature is dominated by studies that have required decades of field data collection to generate sufficiently deep pedigrees. Finally, some of the potential applications of genomic prediction in wild populations are discussed.

## Introduction

A major aim of evolutionary quantitative genetics is to measure and understand heritable genetic variation, and to explain the maintenance of that variation in the face of natural and sexual selection and genetic drift (Walsh and Lynch, 2018). One way that researchers have tried to answer these questions is by conducting long-term ecological studies of natural populations (Charmantier et al., 2014, Kruuk et al., 2008). There are now several studies, mostly of vertebrates, where individual life-histories of entire cohorts have been collected for several decades, spanning 10s of generations of the focal organism. Typically, the pedigree of the population has been determined, allowing researchers to use either (i) quantitative genetics (Kruuk, 2004) and/or (ii) gene mapping approaches (Slate et al., 2010) to study how selection and evolution have shaped diversity. Both approaches have led to genuine breakthroughs in our understanding of how genetic variation and selection have combined to shape biodiversity, but they also both have limitations (especially when applied to natural populations). Proponents of both approaches are aware of these limitations, and they are actively seeking solutions (Charmantier et al., 2014).

The most obvious limitations of a quantitative genetics approach are that (i) the loci explaining trait variation cannot be identified, and (ii) that, in the absence of experimental crosses outside of the natural environment, several generations of data have to be collected before genetic analysis is possible. Gene mapping-based solutions to understanding heritable genetic variation also suffer from problems. Most quantitative trait loci (QTL) identified by linkage mapping in wild populations do not reach strict genome-wide statistical significance, they have not been fine mapped with any precision (i.e. confidence intervals span 10s of centiMorgans), and the effect sizes of reported QTL are almost certainly substantially overestimated (Slate, 2013, Slate et al., 2010). The arrival of next generation sequencing and related technologies such as SNP chips made it possible to type large numbers of individuals at very many (1000s to hundreds of thousands) of SNPs and move from linkage mapping to genome-wide association studies (GWASs). GWAS studies of wild populations have had some notable successes, although again, usually only when large effect loci are segregating (Johnston et al., 2011, Comeault et al., 2014). The gene mapping approach, whether conducted by linkage mapping or by GWAS, has been successful at identifying genes that underlie simple Mendelian traits, but less so for quantitative traits. QTL and GWAS studies require very high statistical significance thresholds (P < 1 × 10^−7^ is not uncommon) to distinguish true positives from false ones. As a result, mapping studies are biased towards detecting causal loci of large effect, because only large effect loci will reach genome-wide significance. If a trait is polygenic then much of the genetic variation will remain unmapped. The most (in) famous example of this is human height, where in a sample size of ∼¼ million people, 697 genomewide significant loci only explained 16% of the heritability (Wood et al., 2014). Studies of natural populations typically involve sample sizes of hundreds, or at most, a few thousand individuals, and therefore statistical power to detect loci of medium-small effect is low. This has been recently demonstrated in collared flycatchers, where association mapping with SNP chips and whole-genome resequencing of individuals with extreme phenotypes failed to identify loci that explained variation in the iconic sexually-selected trait, male forehead patch size (Kardos et al., 2016). The problem may be exacerbated if phenotypes are hard to measure in the field or if environmental heterogeneity is a further source of phenotypic variation. Put simply, even in very large experiments, the power to detect most causal genes is very low. In natural populations, where sample sizes are currently smaller than in studies of human or model organisms, the problem is far greater. This makes it impossible to describe trait architectures or to understand the relationship between genotypes and phenotypes. How then, can evolutionary quantitative geneticists use genomics tools to understand and predict the genetic architecture and microevolution of their focal traits?

One possible solution to the low power / high polygenicity problem is to switch focus away from the hunt for individual significant loci, and instead try to understand the genetic component of phenotypic variation by using information from all of the typed SNPs. Two conceptually different approaches can be taken. The first involves running an ‘animal model’ with the genetic relationship matrix estimated from markers rather than a pedigree. It was recognised several decades ago that allele-sharing at markers could be used to estimate relatedness between pairs of individuals, and that provided there was variance in relatedness, the relatedness coefficients could then be regressed on phenotypic similarity to predict heritabilities (Ritland, 1996, Ritland and Ritland, 1996). Unfortunately, the reliable estimation of relatedness, especially between distant relatives, requires several orders of magnitude more markers than could realistically be typed at that point. As genotyping technology improved, methods using marker-based relatedness became more tractable. In fact, marker-based approaches are potentially more powerful than pedigree-based methods because the realised relatedness between individuals rather than the expected relatedness can be estimated e.g. (Visscher et al., 2006, Hayes et al., 2009b). Using a genomic relationship matrix (GRM) estimated from marker data, instead of the pedigree-derived A matrix, it is possible to estimate quantitative genetic parameters in a traditional mixed model Best Linear Unbiased Predictor (BLUP) framework (VanRaden, 2008). These models have been used to estimate trait heritabilities in natural populations (Robinson et al., 2013, Santure et al., 2013, Bérénos et al., 2014, Silva et al., 2017) as well as in humans (Yang et al., 2010) and agriculturally important organisms (VanRaden et al., 2009). An extension of the model, where the GRM is estimated from a subset of markers, has been used to partition genetic variance into specific parts of the genome, e.g. in chromosome-partitioning (Visscher et al., 2007) or regional heritability mapping (Nagamine et al., 2012) studies.

In addition to providing estimates of parameters such as a trait’s heritability, individual breeding values can also be estimated from genomic best linear unbiased prediction (GBLUP) models. For a historical background and accessible summary of these developments see Hill (2014). An assumption of the GBLUP approach is that the distribution of all of the marker effects are sampled from a single normal distribution; in other words all markers explain some of the variation and the trait is highly polygenic. Of course, this assumption is unrealistic – many quantitative traits are highly polygenic, but not all of the markers will contribute to phenotypic variation. Furthermore, traits with non-polygenic architectures are frequently of interest to breeders, medical geneticists and evolutionary geneticists studying natural populations. A potential improvement on GBLUP methods then, is to use methods that can accommodate different genetic architectures. This is achievable with a second major approach, whereby marker effects are estimated, and can come from more than one (not necessarily normal) distribution. The approach was first described almost twenty years ago (Meuwissen et al., 2001), and has since revolutionised animal and plant breeding and has applications in human personalised medicine (de los Campos et al., 2013b). The concept is deceptively simple. Usually two steps are involved – marker effect sizes are estimated and used to predicted breeding values, and then the accuracy of the breeding values is estimated before any application in a breeding program. In a first stage, marker effects are estimated, usually by Bayesian MCMC methods, in a ‘training population’ of individuals with known phenotypes that are typed at many markers. Next, another panel of individuals with unknown phenotypes (the ‘validation’ or ‘test’ population) is genotyped at the same markers, and their genomic estimated breeding values (GEBVs) are predicted from their genotypes and the estimated effect sizes of each marker. In principle, even in samples containing only distant relatives, the approach works, provided that the markers are in sufficiently strong linkage disequilibrium (LD) with the unknown loci that cause trait variation. If marker density is sufficiently high that adjacent markers are in LD with one another, then they should also be in LD with unknown causal loci. All of the approaches for estimating genomic breeding values from phenotypes and marker data have collectively become known as genomic prediction (and their application in breeding programs, as genomic selection).

Meuwissen and colleagues (2001) described two Bayesian genomic prediction models, Bayes A and Bayes B. In Bayes A, the marker effects are all non-zero, but are drawn from a scaled-inverse χ^2^ distribution, allowing some markers to have large effects. In Bayes B, there are two distributions of marker effects, with a proportion of markers assumed to have zero effect and the remainder with effects drawn from a scaled-inverse χ^2^ distribution. Following the development of Bayes A and Bayes B, further refinements, such as the marker effects coming from different types of distribution, have been made (Gianola, 2013, Habier et al., 2013), but the main principal of using a training population to estimate marker effects, which are then used to predict phenotypes in a genotyped test population applies.

The accuracy of genomic prediction can be estimated by correlation between GEBV and phenotype, divided by the square root of the heritability (Meuwissen et al., 2013). Accuracy depends on a number of factors such as the heritability (*h*^*2*^) of the trait (Daetwyler et al., 2008, Goddard, 2009, Hayes et al., 2009a), the training population sample size (Habier et al., 2013, Meuwissen et al., 2001), the marker density (Habier et al., 2009, Meuwissen and Goddard, 2010), the underlying genetic architecture (i.e. number and effect size of loci) of the trait (Daetwyler et al., 2010) and the statistical approach used (Crossa et al., 2010, Daetwyler et al., 2013, de los Campos et al., 2013a, Meuwissen et al., 2001). Empirical investigation of these factors on genomic prediction accuracy have been described in the animal and plant breeding literature (Heffner et al., 2011, Zhao et al., 2012, Hayes et al., 2010, Hayes et al., 2009a). Highly heritable traits generally have more accurate GEBVs because the environmental and non-additive genetic effects are relatively small, meaning marker effects are more reliably estimated. Increasing the size of the training population allows for more accurate estimate of marker effects; consequently, prediction accuracy is also improved. Accuracy is generally greater with a higher marker density, because markers will be in stronger linkage disequilibrium with unknown causal loci. Conversely, accuracy tends to be lower in populations with larger effective population sizes, because linkage disequilibrium between markers and causal loci is usually weaker. More formal descriptions of how these factors affect accuracy are available elsewhere (Daetwyler et al., 2008, Goddard, 2009, Meuwissen et al., 2013, Hayes et al., 2009a).

The effects of trait genetic architecture and the method employed for the genomic prediction are connected. Because the true genetic architecture, i.e. the actual distribution of effect sizes is generally unknown, it is important that the posterior distribution of marker effects matches the true (but unknown) distribution of SNP effects as closely as possible. Methods where all markers have an effect drawn from a single normal distribution are unlikely to be good predictors of phenotypes determined by a small number of large effect loci. Thus, methods that are flexible enough to capture different genetic architectures are potentially the most useful (Daetwyler et al., 2013). For example, methods that model marker effects as a mixture of different distributions can accommodate large and small effect loci (Erbe et al., 2012, Zhou et al., 2013).

While genomic prediction has become a widely used tool in animal and plant breeding, it has yet to be widely adopted in quantitative evolutionary genetics studies of wild populations, even though there are a number of long-term longitudinal studies where suitable phenotypic and genomic data have been collected. Therefore, the main motivation of this study was to investigate the feasibility and accuracy of genomic prediction in Soay sheep. Previous genetic studies of Soay sheep (Bérénos et al., 2015, Gratten et al., 2007, Gratten et al., 2010, Johnston et al., 2011) have demonstrated that some recorded phenotypes have a simple Mendelian basis (e.g. coat colour, coat pattern), others are highly polygenic (e.g. adult weight), and some are intermediate with many genes of small effect and a small number of loci with quite large effect contributing to genetic variation (e.g. horn length). Thus, a second aim of the study is to assess the accuracy of genomic prediction in different types of trait. Finally, the third aim of the study is to compare the accuracy of different genomic prediction methods for traits with different architectures (Daetwyler et al., 2013, Meuwissen et al., 2013, de los Campos et al., 2013a). Thus, we compare five different methods: (i) GBLUP, (ii) BayesA & (iii) BayesB – the models first introduced by (Meuwissen et al., 2001), (iv) Bayesian Lasso – a model similar to GBLUP and BayesA in the sense that it assumes all SNPs explain some variance and (v) BayesR – a model that assumes SNPs either have zero effect or come from a mixture of >1 Gaussian distribution, potentially with some large effect loci.

## Materials and methods

### Study population

The Soay sheep is a primitive feral breed that resides on the St Kilda archipelago, off the NW of Scotland. Since 1985, the population on the largest island, Hirta (57°48’ N, 8°37’ W), has been the subject of a long-term individual based study (Clutton-Brock et al., 2004). Briefly, the majority of the sheep resident in the village Bay area of Hirta are ear-tagged and weighed shortly after birth and followed throughout their lifetime. Ear punches and blood samples suitable for DNA analysis are collected at tagging. Every year during August, adult sheep and lambs are captured and morphological measurements are taken. Winter mortality is monitored, with the peak of mortality occurring at the end of winter/early spring, and ca. 80% of all deceased sheep are found. To date, extensive life history data have been collected for over 10,000 sheep. More details of the study can be found elsewhere (Clutton-Brock et al., 2004).

### Genetic data

Genotyping of the population was performed using the Illumina Ovine SNP50 beadchip array, developed by the International Sheep Genomics Consortium (ISGC) (Kijas et al., 2012). Genotyping was performed at the Wellcome Trust Clinical Research Facility Genetics Core (Edinburgh, UK). Details about the genotype calling and quality control of the data are available elsewhere (Bérénos et al., 2015, Bérénos et al., 2014, Johnston et al., 2016, Johnston et al., 2011). Briefly, pruning of SNPs and individuals was performed using Plink v 1.9 (Chang et al., 2015). SNPs were retained if they had a minor allele frequency of at least 0.01, a call rate of at least 0.98 and a Hardy-Weinberg Equilibrium Test p value > 0.00001. Individuals were retained if they were typed for at least 0.95 of SNPs. Only autosomal SNPs were analysed, as most genomic prediction software cannot distinguish between autosomal and sex-linked loci; the sheep X chromosome represents ∼5% of the total genome. Because SNP pruning was performed on a per-trait basis, the proportion of SNPs retained after pruning varied slightly between traits (fewest SNPs = 35,885 for coat colour and coat pattern; most SNPs = 36,437 for horn length).

### Phenotypic data

We studied phenotypic data collected for eight different traits: foreleg length, hindleg length, weight, metacarpal length, jaw length, horn length (normal-horned males only), coat colour and coat pattern. An aim of the study was to investigate whether comparisons between the alternative genomic prediction methods were sensitive to the underlying genetic architecture. Therefore, we chose to study traits that had been the focus of previous gene mapping studies and were known to have variable architectures (summarised in Table 1). Among the body size traits, foreleg length, hindleg length, weight and horn length were measured on live animals, whereas metacarpal length and jaw length were taken from cleaned post-mortem skeletal material. Coat colour and coat pattern are independent discrete traits that are recorded at the time of capture and remain fixed throughout life (Clutton-Brock et al., 2004, Gratten et al., 2007, Gratten et al., 2010). Further details of how the morphological traits are measured can be found elsewhere (Bérénos et al., 2015, Bérénos et al., 2014, Johnston et al., 2011). Analyses of live-capture morphological data were restricted to animals captured in August, that were at least 28 months old, to remove most complications of growth and give consistency with previous genetic studies of these traits (Bérénos et al., 2015, Bérénos et al., 2014). Male horn length included measurements taken outside of August, as many males are captured and measured during the rut in November. Post-mortem measurements were likewise restricted to animals who were at least 28 months old when they died.

**Table 1:** Summary of sample sizes and genetic architecture of the eight morphological traits

| Trait | N | Architecture | Important loci/genes | Previous heritability | Heritability* (this study) | Reference |
| --- | --- | --- | --- | --- | --- | --- |
| **Weight** | 1168 | Polygenic | - | 0.30 | 0.32 (0.23-0.43) | (Bérénos et al., 2015, Bérénos et al., 2014) |
| **Jaw length** | 897 | Polygenic | - | 0.50 | 0.59 (0.44-0.71) | (Bérénos et al., 2015, Bérénos et al., 2014) |
| **Foreleg length** | 1126 | Polygenic | Chr16: s23172.,<br>Chr19: s74894.1 | 0.25 | 0.53 (0.43-0.64) | (Bérénos et al., 2015, Bérénos et al., 2014) |
| **Hindleg length** | 1139 | Polygenic | Chr16: s23172.1 | 0.45 | 0.50 (0.38-0.59) | (Bérénos et al., 2015, Bérénos et al., 2014) |
| **Metacarpal length** | 890 | Polygenic | Chr16: s23172.1,<br>Chr19: s74894.1 | 0.50 | 0.62 (0.52-0.75) | (Bérénos et al., 2015, Bérénos et al., 2014) |
| **Male Horn length** | 472 | Oligogenic | Relaxin-like receptor 2 (RXFP2) | 0.30 | 0.42 (0.24-0.58) | (Johnston et al., 2011, Johnston et al., 2013) |
| **Coat colour** | 4737 | Mendelian (dark coat caused by dominant allele) | Tyrosinase-related protein 1 (TYRP1) locus | - | - | (Gratten et al., 2007) |
| **Coat pattern** | 4737 | Mendelian (wild-type coat caused by dominant allele) | Agouti signalling protein (ASIP) | - | - | (Gratten et al., 2010) |

### Modelling non-genetic effects

The quantitative traits being investigated are known to vary between sexes and age classes and are influenced by environmental effects (Bérénos et al., 2014). Because some of the software being tested does not allow the inclusion of repeated measurements or fitting of random effects, it was necessary to fit models with non-genetic effects prior to performing the genomic prediction. Therefore, mixed effects models were run that adjusted phenotypes for fixed effects (sex and age of the animal) and non-genetic random effects (birth year, capture year and animal identity). Traits recorded on live animals had repeated measurements, so individual identity was fitted to remove any permanent environmental effect on phenotype. Traits recorded on skeletal material were recorded just once per animal. Models were run using the *lmer* function in the R v3.6.0 package lme4 v1.1-23 (Bates et al., 2015). The random effect of individual identity was extracted from each model and retained as the phenotype to be analysed in the genomic prediction models. The two Mendelian traits are categorical traits that are fixed throughout lifetime, unaffected by environmental conditions and with the same penetrance in each sex; therefore no phenotypic adjustments were necessary. Details of the model outputs are summarised in Supplementary Table 1.

### Running Genomic Prediction models

A major aim of this study was to compare different methods for obtaining genomic estimated breeding values (GEBVs). Below we briefly discuss these methods, which rely on subtly different assumptions about the distribution of marker effect sizes, and outline how each model was run.

#### BayesA

The BayesA method was one of the two models in the first genomic prediction paper (Meuwissen 2001). All markers have a non-zero effect size, and the marginal distribution of marker effects follows a t-distribution. Here, BayesA was implemented in the R package BGLR v1.0.8 (Perez and de los Campos, 2014). The analyses were performed with default parameter settings except the number of iterations was set to 120,000 (default = 5,000) with a burnin of 20,000 (default= 1,000) and a thinning interval of 100 (default =10).

#### BayesB

The BayesB method of genomic prediction was also introduced in Meuwissen et al, 2001. In this model, the assumption that all markers have a non-zero effect is removed. Instead, the BayesB method assumes a mixture of two distributions, such that some markers have zero effect and others, following a t-distribution, have non-zero effects, with some potentially of large effect. In reality, many loci explain zero genetic variance. BayesB analyses were also performed in the BGLR package with default parameter settings except the number of iterations, burn-in length and thinning interval, which were the same values as used in the BayesA analyses.

#### Bayesian Lasso (BayesL)

This method of genomic prediction is similar to BayesA in that all SNP effects come from a single distribution. However, BayesL models marker effects as following a double exponential distribution rather than a t distribution. A consequence of using this distribution is that, relative to the BayesA method, SNPs with small effect sizes are more strongly shrunk and SNPs with larger effect sizes are less strongly shrunk (de los Campos et al., 2009). BayesL was also implemented in the R package BGLR, using default parameters except the number of iterations, burn-in length and thinning interval, which were the same values as for the BayesA and BayesB analyses.

#### GBLUP

This method assumes that marker effects are drawn from a normal distribution. Rather than estimate the effect of each SNP, GBLUP utilises the genomic relationship matrix (GRM) which is estimated from the proportion of alleles that are identical-by-state (IBS) between every pair of individuals. Here, the GRM was calculated from all typed markers. Genomic Best Linear Unbiased Predictions (GBLUPs) of breeding values are obtained by examining the covariance between phenotypic similarity and pairwise relatedness. GRMs have previously been used to estimate the heritability of traits in the Soay sheep population (Bérénos et al., 2015, Bérénos et al., 2014), but not to obtain GEBVs. As with the BayesA, BayesB and BayesL analyses, BGLR was used to obtain genomic EBVs.

#### BayesR

BayesR models SNP effects as a mixture of >1 normal distribution and an additional component of zero variance (Erbe et al. 2012). We implemented the method within the BayesR v1 software package (Moser et al., 2015) and used the default priors of 4 mixture components with variances of 0, 0.0001, 0.001 and 0.01 of the phenotypic variance. Dirichlet priors for the number of pseudo-observations (SNPs) in each distribution were set to 1, 1, 1 and 5. Priors for the genetic and residual variances were chosen as a scaled inverse-chi squared distribution with scaling parameter estimated from previous pedigree-based animal models (see references in Table 1) and degrees of freedom set to 10. Note that the default parameters give very similar values for genomic estimated breeding values, but we find that some samples of the MCMC chain return zero or very low estimates of additive genetic and residual variance under the default settings (see Supplementary Material Table S2). As with all analyses run in BGLR, the BayesR analyses were run for a total of 120,000 iterations with a burnin of 20,000 and a thinning interval of 10.

Because BayesR reports the genetic variance explained by each SNP, SNP effect sizes can be aggregated (e.g. across the genome, across individual chromosomes) to describe the genetic architecture of the trait. Detailed descriptions of the trait architectures are not the main goal of this work, but we do report them in the Supplementary material. This is partially motivated by the knowledge that many of our focal traits already have well-described architectures (Table and so it is useful to examine whether BayesR is assigning amounts of variance to loci that are consistent with genome wide association studies. We also compare descriptions of trait architectures between BayesR models using the default parameters and models using priors informed by previous quantitative genetic studies of those traits (Table S2).

### Assessing prediction accuracy and bias

Genomic prediction accuracy was assessed by cross-validation; each data set was randomly split into a training population containing 95%, 50% or 10% of individuals and a test population containing the remaining 5%, 50% or 90% of individuals. Twenty replicates were generated for every combination of trait, model and training population size (8 traits * 5 methods * 3 training sizes * 20 replicates = 2400 models). For the six continuous traits, prediction accuracy was estimated as the correlation between the GEBV and an individual’s phenotype, divided by the square root of the heritability (i.e. *r*_GEBV,y/_*h*, where h^2^ is the heritability and *y* is the phenotype) Accuracy was averaged across the 20 replicates. The phenotypes were the values pre-adjusted for age, sex, year of birth and year of death using the same models as described in the section on *Modelling non-genetic effects.* Because the phenotypes were adjusted for random and fixed effects, the heritability of each trait was estimated from the adjusted phenotypes. These estimates of heritability will be greater than previously published estimates, because non-genetic sources of variation have already been removed, but they are more appropriate for measuring the observed and expected accuracy of the GEBVs. Trait heritabilities were estimated using Animal models run in the R package MCMCglmm v2.29 (Hadfield, 2010). In these models the additive genetic variance was estimated from a relationship matrix determined by the population pedigree; the pedigree was constructed from a combination of behavioural observations and genotype data from 315 SNPs. Pedigree construction methods are described elsewhere (Bérénos et al., 2014). MCMCglmm models were run for 120,000 iterations with a burn-in of 20,000 and sampling every 100^th^ iteration. In addition to recording the correlation between GEBVs and phenotypes, a linear regression was performed to estimate the slope between the two variables. A slope of 1 is indicative of there being no bias in GEBVs (Meuwissen et al., 2001, Moser et al., 2015). Slopes greater than one would suggest that between-individual differences in GEBVs underestimate the difference in phenotypic values.

For the two categorical traits (Coat colour/pattern), accuracy was determined by measuring the area under the curve (AUC) in a Receiver Operating Characteristic (ROC) curve. ROC curves are well established tools for assessing how well genetic markers predict categorical traits such as disease presence/absence (Wray et al., 2010). ROC curve analysis were performed with the R package pROC v1.16.2 (Robin et al., 2011).

## Results and Discussion

Attempts to use genomic prediction in an ecological context remain very rare compared to its application in breeding or medicine, although some examples are now described (Stocks et al., 2019, Gienapp et al., 2019, Bosse et al., 2017). As far as we are aware this is the first genomic prediction study in a wild population that considers either multiple traits with different genetic architectures or different genomic prediction methods.

### Genomic Prediction Accuracy

The main finding is that genomic prediction in the Soay sheep population is accurate regardless of trait architecture or of the method used (Figure 1, Table S2), even when training sizes are relatively modest (a few hundred individuals). For the continuous traits, accuracy was typically high (∼0.70) when most of the phenotyped animals were included in the training set (Table S2, Figure 1a). Unsurprisingly, the accuracy declines when the training size is decreased. The accuracy of jaw length GEBVs were a little lower than for the other traits. It is not immediately obvious why jaw length would have a lower accuracy, as it is not less heritable than the other traits, and although it is highly polygenic – no QTL have previously been identified in GWAS studies – it is no different to adult weight in that regard (Bérénos et al., 2015). The phenotypes of the two categorical traits were predicted with very high accuracy, even when only 10% of animals were used in the training dataset (Table S2, Figure 1b). As expected, for all traits, the accuracy was generally lower when the training set was smaller (Table S2, Figure 1).

**Figure 1:**
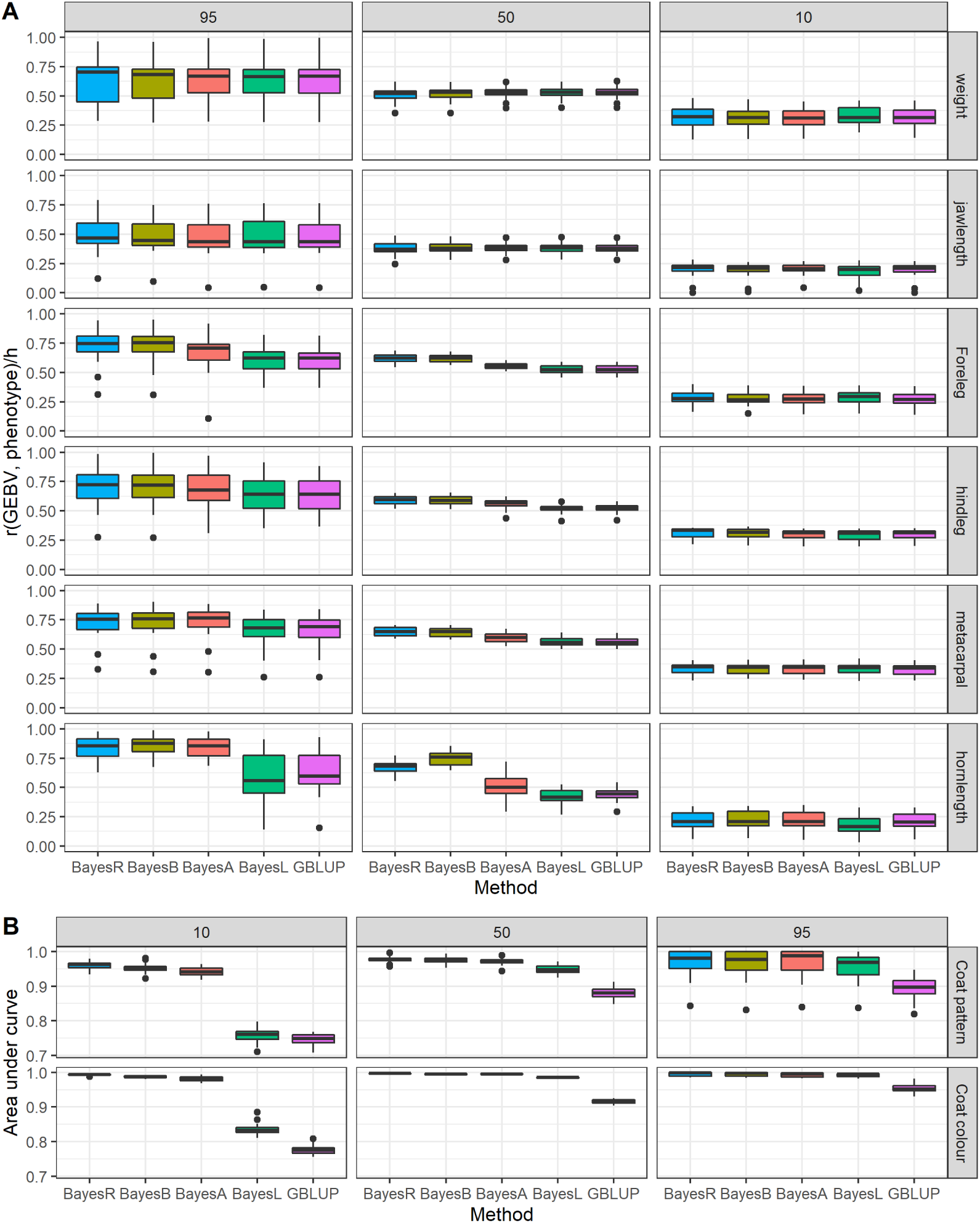
Accuracy of genomic prediction for (a) quantitative and (b) Mendelian traits. For quantitative traits, accuracy is measured as r_GEBV,y_/h, the mean correlation between GEBVs and the phenotype, divided by the square root of the heritability. For Mendelian traits, accuracy is determined as Area Under Curve from ROC analysis. All plots show the mean and SE obtained from 20 replicates of training sizes comprising 95%, 50% or 10% of the available individuals.

Comparisons between studies are not straightforward because accuracy depends on both biological (e.g. genome size, effective population size, heritability) and technical (e.g. marker density, training population size) factors (Daetwyler et al., 2010, Daetwyler et al., 2008, Goddard, 2009, de los Campos et al., 2013b), which inevitably will vary between studies. Certainly, the accuracy reported here is comparable to situations in plant and animal breeding where genomic selection is routinely used for trait improvement (Lin et al., 2014, Hayes et al., 2009a) and is probably greater than has been observed for morphological traits in commercial sheep populations (Auvray et al., 2014, Brito et al., 2017). Therefore, the prospects for genomic prediction in this population are promising.

### Genomic prediction bias

If genomic prediction estimates of GEBVs are unbiased, then it is expected that regressions of GEBVs on phenotypes should be equal to 1.0 (Meuwissen et al., 2001).There was a tendency for regression coefficients to be a little less than 1.0, meaning GEBVs overestimated between-individual variation, but this was largely driven by the models with smallest training sizes (Figure 2, Table S3). In the 50% and 95% training population datasets, there was relatively little bias with regression coefficients close to 1.0 for most traits and most models. Taking an average across 360 analyses (20 replicates x 3 training sizes x 6 quantitative traits) BayesL had the mean regression coefficient closest to 1.0 (Table S3, top row). However, of all of the methods, it had by far the largest standard deviation of regression coefficient, and therefore may be the most prone to severe bias. Of the remaining methods, BayesR had the mean regression coefficient closest to 1.0 and with the lowest standard deviation, although all of the other methods performed quite similarly with respect to bias (Table S3).

**Figure 2.**
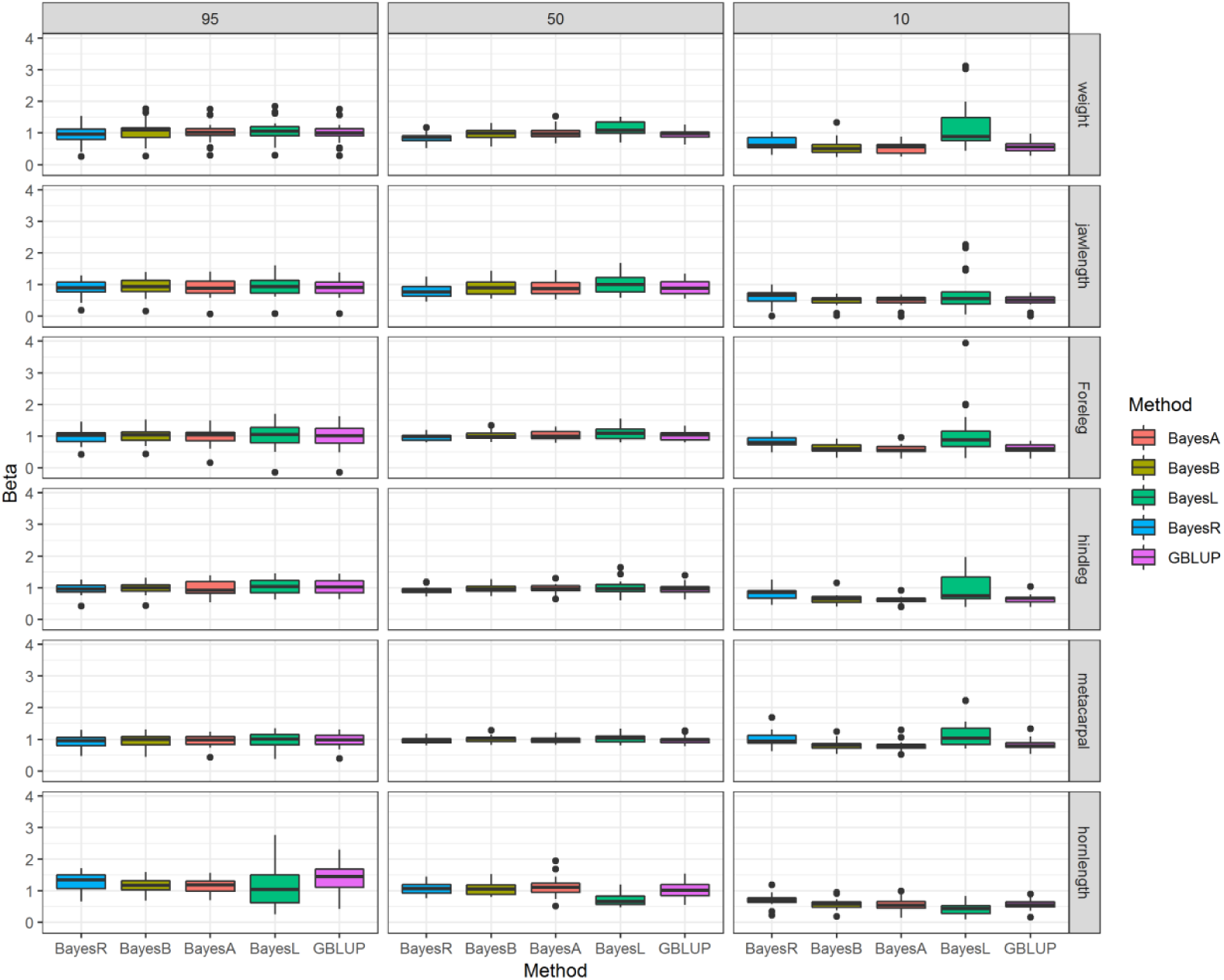
Boxplots showing the mean and distribution of regression coefficients when GEBVs were regressed on phenotypes for the 6 continuous traits. Each boxplot summarises 20 replicates per trait/training size/method combination.

### Comparison of Models

Among most of the continuous traits, there was little difference in the accuracy (Table S2, Figure 1a). This is not surprising as all of the models either make an assumption that the trait is polygenic, or can explicitly model the trait as polygenic. BayesR and BayesB were perhaps marginally more accurate than BayesA, BayesL and GBLUP. For horn length, the most oligogenic of the continuous traits, and the two single-locus Mendelian traits, coat colour and coat pattern, BayesR and BayesB were the most accurate (Table S2, Figure 1b), presumably because they allow for the modelling of both zero effect and major effect loci. BayesA, was the next best performing model, with BayesL and GBLUP being considerably less accurate than all others, especially with smaller training populations. Thus, across all types of trait architecture, and considering both accuracy and bias, BayesR and BayesB appeared to be the most reliable methods of those examined here. Previous studies of humans and livestock have reached similar conclusions (Moser et al., 2015, Erbe et al., 2012). From this point, we mostly focus on results from the BayesR analyses, as that is the most accurate approach, and because it allows a more detailed investigation of trait genetic architecture.

### Comparisons between observed and expected accuracy

The relatively high accuracy of genomic prediction in Soay sheep is perhaps not especially surprising. It is possible to predict the accuracy of genomic prediction if information about the effective population size, genome size, recombination rate, trait heritability, and the genomic architecture are available (Daetwyler et al., 2010, Goddard et al., 2010). Here we used equation 2 of Daetwyler et al (2010):

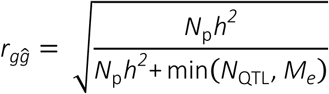

where *r*_*gĝ*_ is the correlation between the true breeding value *g* and the GEBV *ĝ*. *r*_*gĝ*_ is equivalent to the *r*_GEBV,y_/*h*, the measure of accuracy we use in this paper (Meuwissen et al., 2013). *N*_p_ is the number of individuals in the training set, *h*^2^ is the trait heritability and *N*_QTL_ is the number of independent loci contributing to the trait. The other parameter, *M*_*e*,_ is the estimate for the number of independent chromosomal segments, given by *M*_*e*_ = 2*N*_*e*_*L*/log(4*N*_*e*_*L*) where *N*_*e*_ is the effective population size and *L* is the genome linkage map length, measured in Morgans. The length of the Soay sheep genome is 33.04 Morgans (Johnston et al., 2016), and the effective population size is around 194 – see the supplementary material of Kijas et al. (2012) – giving an *M*_*e*_ of 1263. We used the pedigree-estimated heritabilities of the traits adjusted for fixed and random effects (Table 1). We estimated the number of causal loci as 3,000 (weight and jaw length), 1,000 (foreleg, hindleg, metacarpal), 100 (horn length) or 1 (coat colour and coat pattern). 100 loci may seem like a large number of loci for horn length, given one locus explains most of the additive genetic variance (Johnston et al., 2013, Johnston et al., 2011), but the BayesR analysis suggests that the remaining additive genetic variation is likely to be determined by many loci (Table S4). Note that while the true number of QTL can only be crudely guessed, the predicted accuracy is a function of whichever is smaller of *N*_*QTL*_ and *M*_*e*_; it is likely that for polygenic traits of Soay sheep *M*_*e*_ is lower than *N*_*QTL*_, making the expected accuracy insensitive to guesses of the true number of QTL. Overall, for all three training population sizes (95%, 50% and 10%), there was a good fit between the observed and expected accuracy of genomic prediction (Fig 3); for the leg length and weight traits, the observed accuracy was a little higher than expected.

**Figure 3:**
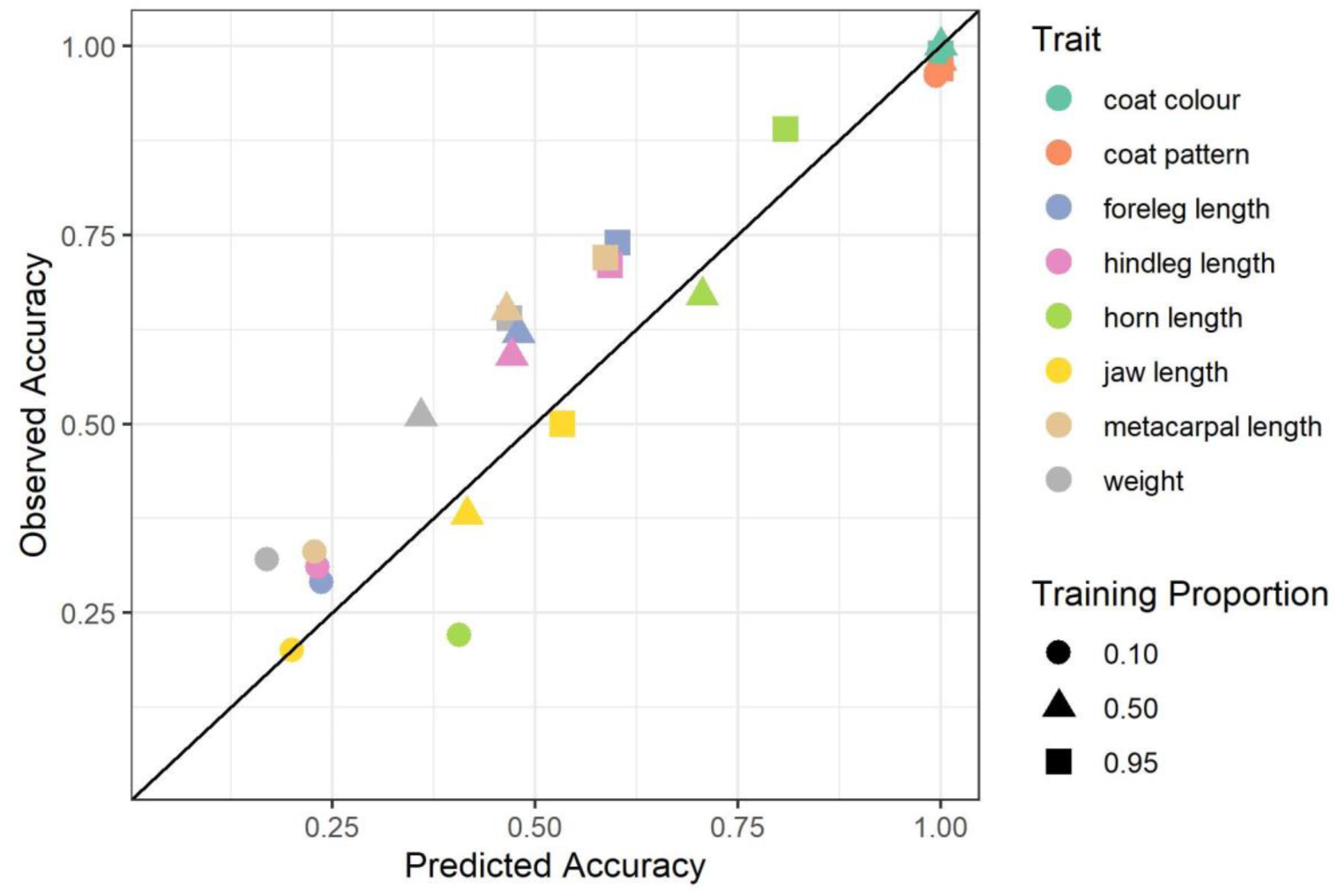
Observed and Predicted Accuracy of BayesR genomic prediction in Soay sheep

The reasons for the accuracy of some traits being slightly better than expected are unclear. One possible explanation is that the effective population size has been overestimated, which would result in a lower predicted accuracy than if the correct *N*_*e*_ had been used. However, overestimating *N*_*e*_ would be expected to affect the predicted accuracy of all traits, not just a subset of them. It is notable that previous GWAS studies (Bérénos et al., 2015) of the three leg traits identified QTL of reasonably large effect (up to 15% of the additive genetic variance), despite the traits being reasonably polygenic. These findings are supported by the BayesR analyses of genetic architecture (see Table S2). In general the traits with larger effect size loci had the greatest accuracy, so perhaps the greater than expected accuracy of leg length and weight traits is attributable, at least in part, to these relatively large effect loci.

### Description of trait architectures by BayesR

Although a description of trait architectures was not a primary objective of this work, BayesR provides estimates of many parameters of interest, including the trait heritability and the number and effect sizes of SNPs contributing to phenotypic variation. We summarise the trait architectures in the supplementary material. Because most of the traits considered here have been the focus of previous studies, it is possible to compare the BayesR outputs with other approaches such as GWAS and chromosome partitioning analyses. Trait architectures of the eight traits are summarised in Table S4, chromosome partitioning plots are presented in Fig S1, and Manhattan plots of posterior inclusion probabilities of each SNP having a non-zero effect size are presented in Fig S2. Broadly, the genetic architectures described by BayesR are consistent with those reported previously. Estimates of trait heritability were similar to estimates made using pedigrees (for comparisons see Tables 1 and S4), with the six continuous traits all having BayesR estimated heritabilities between 0.39 and 0.64. Coat colour and coat pattern had lower heritabilities, despite both being determined by a single locus; this is expected as both *TYRP1* (coat colour) and *ASIP* (coat pattern) have one allele that is completely dominant to the other, so some non-additive (dominance) genetic variance at the causal loci is well documented (Gratten et al., 2007, Gratten et al., 2010).

BayesR returns an estimate of the number of SNPs that explain the phenotypic variation. We urge caution against taking these estimates as definitive (and note that the 95% confidence intervals span a ∼10-fold range for most traits; Table S4). However, the estimated number of SNPs was greatest for the traits that were thought to be highly polygenic (weight, jaw length), and was least for the Mendelian (coat colour and coat pattern) and major locus (horn length) quantitative traits. The estimated number of causal SNPs was intermediate for those quantitative traits where moderate effect size QTL had been discovered in previous GWAS studies. Thus, within-population, between-trait comparisons in the number of SNPs may give an indication of which traits are more/less polygenic, provided the same set of markers is used for each trait.

Chromosome partitioning plots (Fig S1) were broadly consistent with previous work using chromosome-wide relatedness matrices (Fig 1 of Bérénos et al., 2015). However, in that study, two traits (adult weight and jaw length) had significant correlations between chromosome length and the proportion of variation explained by the chromosome, but in this study neither trait showed a significant relationship. Nonetheless, the chromosomes that tended to explain the most variation in the earlier study, also explained the most variation here. An exception was for jaw length, where Chromosome 20 explained more than 30% of the additive genetic variance here, but did not in the earlier study. However, no SNP on Chromosome 20 had a posterior inclusion probability > 0.5 (Figure S2b), so there is no compelling evidence of a major locus QTL on that chromosome. Instead, posterior inclusion probabilities plots indicate at least two regions of chromosome 20 contribute to jaw length variation (Fig S2b).

The two Mendelian traits, coat colour and coat pattern, are characterised by one allele being dominant to the other (at *Tyrp1* that dark allele is dominant to the light allele and at *ASIP* the wild allele is dominant to the self allele). Despite neither causal SNP being included on the ovine SNP chip, the distribution of GEBVs for both traits was trimodal (Fig S3), indicating that the linked SNPs successfully distinguished between the three possible genotypes (heterozygotes and the alternative homozygotes). In other words, it was possible to distinguish between phenotypically identical animals that are either heterozygous or homozygous for the dominant allele.

### Comparison between default and informed priors for BayesR models

GEBVs estimated from BayesR models running the default parameters and BayesR models using priors informed by previous molecular genetic studies were almost identical; for all eight traits the correlation in GEBVs between the two models was > 0.996 (Table S2). Genetic architectures were similar between the two models, although there was a tendency for the heritability to be a little greater when using the model that used informed priors (Table S2). For one trait, horn length, the default model tended to have a large autocorrelation between estimates of V_A_ from consecutive samples of the MCMC and it also frequently estimated there to be no additive genetic variance for the trait (almost 10 % of MCMC samples), despite the trait having a high heritability.

### Conclusions and future directions

In this population the accuracy of genomic prediction was comparable to that seen in applied animal and plant breeding programs. In part, this is because Soay sheep have a history of isolation on a small island and a relatively small effective population size. This means LD extends several megabases across much of the genome (Bérénos et al., 2014, Feulner et al., 2013), and the number of SNPs (∼38K) on a medium density chip is sufficient to tag unknown causal loci. Future work will examine whether a higher marker density (400-500K SNPs) results in a further improvement in accuracy. While other study systems used in evolutionary and ecological research may not have LD extending as far as it does in Soay sheep, the era of whole-genome sequencing hundreds or thousands of individuals is upon us. Therefore, an insufficient marker density will soon become a problem of the past, and genomic prediction should be feasible in other populations. Excitingly, this means that quantitative genetic studies are no longer restricted to wild populations with multigenerational pedigrees. This is especially important for long-lived organisms where sampling individuals across several generations may be a decades-long endeavour or for small organisms that are hard to track in the wild such as insects. Instead, cross-sectional rather than vertical sampling of study populations is possible, which greatly expands the range of organisms that could be studied. Genomic methods for describing genetic architectures, like the BayesR approach, mean that many different aspects of genomic architecture (e.g. a trait’s heritability, additive genetic variance, and the contribution of individual loci and individual chromosomes etc.) can be estimated, even in the absence of a pedigree.

Of course, genomic prediction analyses usually seek to estimate breeding values (and by extension, phenotypes) which are properties of the individual rather than of a population (such as, e.g., a trait’s heritability). There are several obvious areas where GEBVs can be used to address long-standing problems in evolutionary ecology. First, being able to estimate an individual’s breeding value and/or phenotype before it is expressed means it should be possible to predict (and test) how different individuals will respond to a future environmental event. A good example of this comes from a genomic prediction study of ash trees experiencing an outbreak of ash dieback, a disease caused by the fungus *Hymenoscyphus fraxineus* (Stocks et al., 2019). Being able to identify which young plants are most resistant to dieback could be an essential tool in re-establishing populations devastated by disease. Second, GEBVs can be used to better understand the ‘invisible fraction’ problem in evolutionary biology, highlighted more than three decades ago (Grafen, 1988). The invisible fraction refers to the idea that the individuals in a population who reach reproducible age, may not be representative of the entire population if mortality up to reproduction is non-random (Hemmings and Evans, 2020, Hadfield, 2008). Because estimating GEBVs in a test population requires only genomic data and no phenotypes, it should be possible to compare GEBVs between all of the individuals born in a cohort and the subset of them that reach reproductive age. Thus, genomic prediction at a fitness or survival-related trait should allow direct testing for, and quantification of, invisible fractions.

Perhaps the most obvious application of genomic prediction in wild populations will be to explore microevolutionary trends and in particular revisit the ‘evolutionary stasis’ problem (Merilä et al., 2001c). This refers to the observation that traits are frequently observed to be heritable and under directional selection, yet they do not always evolve in response to that selection in the expected way. Detailed descriptions and explanations of the possible explanations for stasis are available elsewhere (Kruuk et al., 2001, Merilä et al., 2001b, Merilä et al., 2001a, Merilä et al., 2001c). Quantitative genetic approaches to understand evolutionary stasis have used EBVs derived from animal models applied to pedigreed populations (Wilson et al., 2007, Coltman et al., 2003, Garant et al., 2004), but a number of problems and biases associated with this approach have been highlighted (Hadfield et al., 2010, Postma, 2006). Notably, pedigree-derived predicted breeding values are more strongly correlated with the phenotype than true breeding values are (Postma, 2006). This is especially true for individuals without many phenotyped relatives in the pedigree. This is problematic for studies looking at microevolutionary trends in breeding values, as it means temporal changes in EBVs may reflect environmental effects on phenotypes rather than genetic ones. Similarly, in a pedigree-based approach, those individuals that lack phenotypic records and have few relatives have EBVs very close to the population mean of true breeding values (Hadfield et al., 2010). In studies that explore temporal trends in breeding values, these relatively uninformative individuals may be clustered towards either or both ends of a time series. Both problems are largely overcome with GEBVs generated from a genomic prediction test population, because each individual’s genome rather than its phenotype or number of phenotyped relatives is used to predict its breeding value. It has been shown through simulations that genomic prediction is a more accurate method for estimating EBVs than pedigree-based methods when the focal individual is not closely related to phenotyped individuals (Clark et al., 2012). Of course, other problems highlighted in discussions of using breeding values to study microevolutionary trends, such as failing to incorporate the uncertainty in GEBVs (Hadfield et al., 2010), must still be addressed, but that is relatively straightforward if a Bayesian solution is used. Thus, it seems likely that genomic prediction can pave the way for new analyses of microevolutionary trends.

In conclusion, this study shows that genomic prediction can reliably measure individual breeding values in the Soay sheep population, and that high accuracy does not appear to be restricted to traits with specific underlying genetic architectures. Approaches that can model zero effect, small effect and large effect loci seem to be the most consistently reliable. Finally, we anticipate that similar studies will soon be possible in many other previously under-studied organisms, paving the way towards both applied evolutionary quantitative genetics research and a re-exploration of some classic, yet unresolved, problems.

## Data Availability

Genotype and phenotype data, along with files showing which individuals were allocated to training and test populations can be found on Dryad [data will be posted when manuscript is accepted for publication]

## Author Contributions

Coordination and collection of field data: JGP and JMP. SNP genotype collection and quality control: CB, PAE and SEJ. Data analysis: BA, DCH and JS. Manuscript writing: JS, BA with input from all authors.

## Acknowledgements

We thank the National Trust for Scotland (NTS) for permission to carry out field work on St Kilda and to QinetiQ and Eurest for logistical support. We are grateful to the many volunteers who have helped with field data collection on St Kilda over the last three decades. Genotyping was carried out at the Wellcome Trust Clinical Research Facility Genetics Core. This work was funded by a Natural Environment Research Council (NERC) grant, (NE/M002896/1) awarded to JS and JMP. SNP genotyping was mostly funded by a European Research Council (ERC) grant (Wild Evolutionary Genomics) awarded to JMP.

