## Supplementary Material for "Genomic prediction in the wild: A case study in Soay sheep"

### Supplementary Information: Table of Contents


| | |
|---|---|
| Page 2 | Part 1: Description of mixed models used to adjust phenotypes for non-genetic effects |
| Pages 3-4 | Part 2: Tables summarising the accuracy and bias of genomic prediction |
| Pages 5-9 | Part 3: Tables and Figures summarising the genetic architectures of the eight traits |
| Page 10 | References |

*Part 1: Preadjusting phenotypes for non-genetic sources of variation*

Table S1: Summary of models used to extract age, sex, year and repeated measure adjusted phenotypes. For fixed effects the table shows the coefficient of the term and its standard error. For random effects the amount of variance explained and its standard error are shown. All animals were at least 28 months old at the time of measurement. Weight, foreleg, hindleg and horn length measurements were all collected on live animals, while jaw length and metacarpal length were measured from skeletal material. Age is age at capture for live measurements and age at death for skeletal measurements. Capture year was not fitted for skeletal traits such as Birth Year and Age at death already model that source of variance. The number of animals is slightly larger than the number of individuals going into the genomic prediction models (Table 1) because not all of the phenotyped animals were genotyped. All models were run using the R package lme4.

| | *N. Obs* | *N. animals* | *Fixed Effects* | | *Random Effects* | | | |
|---|---|---|---|---|---|---|---|---|
| *Phenotype* | | | Sex | Age | Birth Year | Capture Year | ID | Residual |
| Weight | 3622 | 1259 | 9.50 (0.18) | 0.33 (0.018) | 0.33 (0.57) | 1.19 (1.09) | 6.13 (2.48) | 3.07 (1.75) |
| Foreleg | 3282 | 1142 | 6.73 (0.34) | 0.12 (0.03) | 0.68 (0.82) | 14.19 (3.77) | 18.92 (4.35) | 8.13 (2.85) |
| Hindleg | 3404 | 1156 | 8.05 (0.43) | 0.07 (0.03) | 2.58 (1.61) | 0.75 (0.87) | 37.82 (6.15) | 6.59 (2.57) |
| Horn length | 987 | 535 | - | 26.36 (0.88) | 335.5 (18.3) | *2133.6 (11.6) | 3221.6 (56.8) | 904.8 (30.1) |
| Jaw length | 965 | 954 | 3.70 (0.34) | 0.76 (0.06) | 0.74 (0.86) | - | 18.80 (4.34) | 0.32 (0.57) |
| Metacarpal | 933 | 922 | 2.78 (0.31) | 0.15 (0.05) | 0.16 (0.40) | - | 14.60 (3.82) | 1.37 (1.17) |

* For male horn length this variance component is for capture month nested within capture year. Capture month was fitted for this trait as animals measured outside of August, primarily during the rut (November) were included in the model and horns may continue to grow between August and November.

#### *Part 2: A summary of the accuracy and bias of the 5 genomic prediction methods*

Table S2: Accuracy of genomic prediction using 5 different methods of estimating GEBVs. Accuracy is reported as the mean (se) correlation between GEBV and the phenotypic value adjusted for non-genetic effects, scaled by the square root of the heritability ($r_{GEBV,y}/h$). For coat colour and coat pattern (discrete traits), the accuracy was assessed by computing AUC in the pROC package. All estimates are derived from 20 cross-validation replicates.

| Training size (%) | BayesR | BayesA | BayesB | BayesL | GBLUP |
|---|---|---|---|---|---|
| Weight (n = 1,168) | | | | | |
| 95 | 0.64 (0.05) | 0.63 (0.04) | 0.64 (0.04) | 0.62 (0.04) | 0.63 (0.04) |
| 50 | 0.51 (0.01) | 0.53 (0.01) | 0.52 (0.01) | 0.53 (0.01) | 0.53 (0.01) |
| 10 | 0.32 (0.02) | 0.31 (0.02) | 0.31 (0.02) | 0.33 (0.02) | 0.32 (0.02) |
| Jaw length (n = 897) | | | | | |
| 95 | 0.50 (0.04) | 0.47 (0.04) | 0.49 (0.04) | 0.47 (0.04) | 0.47 (0.04) |
| 50 | 0.38 (0.01) | 0.38 (0.01) | 0.38 (0.01) | 0.38 (0.01) | 0.38 (0.01) |
| 10 | 0.20 (0.02) | 0.19 (0.02) | 0.19 (0.02) | 0.19 (0.02) | 0.19 (0.02) |
| Foreleg (n =1,126) | | | | | |
| 95 | 0.74 (0.04) | 0.67 (0.04) | 0.74 (0.04) | 0.58 (0.05) | 0.57 (0.05) |
| 50 | 0.62 (0.01) | 0.56 (0.01) | 0.62 (0.01) | 0.53 (0.01) | 0.53 (0.01) |
| 10 | 0.29 (0.01) | 0.28 (0.01) | 0.28 (0.01) | 0.29 (0.01) | 0.28 (0.01) |
| Hindleg (n = 1,139) | | | | | |
| 95 | 0.71 (0.04) | 0.69 (0.04) | 0.71 (0.04) | 0.64 (0.04) | 0.64 (0.04) |
| 50 | 0.59 (0.01) | 0.56 (0.01) | 0.59 (0.01) | 0.52 (0.01) | 0.52 (0.01) |
| 10 | 0.31 (0.01) | 0.30 (0.01) | 0.30 (0.01) | 0.29 (0.01) | 0.30 (0.01) |
| Metacarpal length (n =890) | | | | | |
| 95 | 0.72 (0.03) | 0.73 (0.03) | 0.72 (0.03) | 0.65 (0.03) | 0.65 (0.03) |
| 50 | 0.65 (0.01) | 0.59 (0.01) | 0.64 (0.01) | 0.56 (0.01) | 0.56 (0.01) |
| 10 | 0.33 (0.01) | 0.33 (0.01) | 0.33 (0.01) | 0.33 (0.01) | 0.33 (0.01) |
| Horn length (n = 472) | | | | | |
| 95 | 0.89 (0.03) | 0.91 (0.03) | 0.91 (0.03) | 0.65 (0.05) | 0.61 (0.05) |
| 50 | 0.67 (0.01) | 0.52 (0.02) | 0.75 (0.01) | 0.43 (0.01) | 0.44 (0.01) |
| 10 | 0.22 (0.02) | 0.22 (0.02) | 0.23 (0.02) | 0.17 (0.02) | 0.22 (0.02) |
| Coat Colour (n = 4,737) | | | | | |
| 95 | 0.99 (0.00) | 0.99 (0.00) | 0.99 (0.00) | 0.99 (0.00) | 0.95 (0.00) |
| 50 | 1.00 (0.00) | 0.99 (0.00) | 1.00 (0.00) | 0.98 (0.00) | 0.92 (0.00) |
| 10 | 0.99 (0.00) | 0.98 (0.00) | 0.99 (0.00) | 0.84 (0.00) | 0.78 (0.00) |
| Coat pattern (n = 4,737) | | | | | |
| 95 | 0.97 (0.01) | 0.97 (0.01) | 0.96 (0.01) | 0.95 (0.01) | 0.90 (0.01) |
| 50 | 0.98 (0.00) | 0.97 (0.00) | 0.98 (0.00) | 0.95 (0.00) | 0.88 (0.00) |
| 10 | 0.96 (0.00) | 0.94 (0.00) | 0.95 (0.00) | 0.76 (0.00) | 0.75 (0.00) |

Table S3. Regression of GEBV on phenotype for quantitative traits. Mean (SE) from 20 replicates are reported for each combination of trait, method and training size. Overall mean (SD) is reported for each method in the top row.

| Training size (%) | BayesR | BayesA | BayesB | BayesL | GBLUP |
|---|---|---|---|---|---|
| *Mean (SD)* | *0.90 (0.26)* | *0.87 (0.30)* | *0.87 (0.29)* | *0.99 (0.46)* | *0.88 (0.32)* |
| Weight (n = 1,168) | | | | | |
| 95 | 0.95 (0.08) | 1.03 (0.08) | 1.03 (0.08) | 1.06 (0.09) | 1.02 (0.08) |
| 50 | 0.84 (0.04) | 1.00 (0.04) | 0.97 (0.05) | 1.12 (0.05) | 0.96 (0.04) |
| 10 | 0.67 (0.05) | 0.52 (0.05) | 0.56 (0.04) | 1.23 (0.17) | 0.57 (0.04) |
| Jaw length (n = 897) | | | | | |
| 95 | 0.88 (0.06) | 0.91 (0.07) | 0.93 (0.06) | 0.95 (0.08) | 0.91 (0.07) |
| 50 | 0.79 (0.05) | 0.91(0.06) | 0.93 (0.06) | 1.03 (0.07) | 0.92 (0.05) |
| 10 | 0.60 (0.04) | 0.48 (0.04) | 0.49 (0.04) | 0.77 (0.14) | 0.50 (0.04) |
| Foreleg (n =1,126) | | | | | |
| 95 | 0.99 (0.06) | 0.99 (0.07) | 1.01 (0.06) | 1.01 (0.09) | 0.99 (0.09) |
| 50 | 0.97 (0.03) | 1.03 (0.04) | 1.02 (0.03) | 1.10 (0.05) | 1.02 (0.03) |
| 10 | 0.82 (0.04) | 0.58 (0.03) | 0.62 (0.04) | 1.10 (0.18) | 0.61 (0.03) |
| Hindleg (n = 1,139) | | | | | |
| 95 | 0.95 (0.04) | 0.98 (0.05) | 0.98 (0.05) | 1.03 (0.06) | 1.02 (0.06) |
| 50 | 0.92 (0.03) | 0.97 (0.03) | 0.96 (0.03) | 1.01 (0.05) | 0.97 (0.04) |
| 10 | 0.79 (0.04) | 0.61 (0.03) | 0.65 (0.04) | 0.96 (0.10) | 0.64 (0.03) |
| Metacarpal length (n = 890) | | | | | |
| 95 | 0.94 (0.05) | 0.96 (0.04) | 0.95 (0.05) | 0.98 (0.06) | 0.97 (0.05) |
| 50 | 0.97 (0.02) | 0.98 (0.02) | 1.01 (0.03) | 1.03 (0.03) | 0.98 (0.03) |
| 10 | 1.01 (0.05) | 0.80 (0.04) | 0.81 (0.04) | 1.16 (0.10) | 0.82 (0.04) |
| Horn length (n = 472) | | | | | |
| 95 | 1.28 (0.07) | 1.14 (0.05) | 1.15 (0.06) | 1.18 (0.14) | 1.38 (0.11) |
| 50 | 1.08 (0.05) | 1.12 (0.08) | 1.06 (0.05) | 0.72 (0.05) | 1.02 (0.06) |
| 10 | 0.70 (0.04) | 0.57 (0.04) | 0.58 (0.04) | 0.42 (0.04) | 0.57 (0.04) |

#### *Part 3: A summary of the genetic architecture of the 8 traits, inferred using BayesR*

The BayesR analyses of each trait produce estimates of the effect size of each SNP. This information can be used to make inferences about the genetic architecture of a trait. In this section, the architectures of each trait are described. Note that for these analyses, all phenotyped and genotyped animals are used in the training population i.e. there is no test population.

Table S3 summarises the architectures of each trait, when BayesR was run using the default parameters, and when it was run using priors based on previous estimates of the heritability of each trait. The results were very similar regardless of priors although for one trait, horn length, the MCMC chain sometimes exhibited a strong autocorrelation between different samples of the chain under the default settings, but not when using informed priors.

Figure S1 summarises a chromosome partitioning analysis, where the sum of the SNP effects is calculated for each chromosome.

Figure S2 shows a Manhattan plot, where the probability of each SNP having a non-zero effect is summarised as a posterior inclusion probability, on a 0 to 1 scale. A value near to 1 indicates it is highly likely that the SNP has a non-zero effect size.

Figure S3 shows the distribution of GEBVs for the two Mendelian traits, coat colour and coat pattern. Both distributions are tri-modal, consistent with the previously described one-locus, two-allele models where one allele is dominant to the other (Table 1 of main manuscript). The causal SNPs for coat colour and coat pattern are not on the ovine SNP chip, but clearly their effect on the phenotypes is tagged by SNPs in linkage disequilibrium. The (relatively) small variation around each mode presumably reflects BayesR attributing small amounts of variation to other non-causal loci (see Fig S2).

*Table S4: Description of the genetic architecture of all 8 traits. Results are reported when BayesR was run under the default parameter settings and when using priors informed by previous quantitative genetic studies [1-5]. As a default BayesR uses improper (flat) distributions as priors for $V_A$ and $V_E$. Under the 'informed model' priors for $V_A$ and $V_E$ were specified using the value given in the row Scaling parameter (VA, VE) and assuming 10 degrees of freedom. $V_A$= additive genetic variance; $V_E$ = all other sources of variation; $h^2$ = narrow-sense heritability; N_SNPs = total number of SNPs that explain trait variation; N_SNPs_0.0001, N_SNPs_0.001 and NSNPs_0.01 are the number of SNPs in each of the three non-zero effect size distributions (in order from smallest to largest effect size distribution). PGE_0.0001, PGE_0.001 and PGE_0.01 is the proportion of genetic variance assigned to the three non-zero effect size distributions. Autocorr $V_A$ is the autocorrelation in estimates of $V_A$ reported by consecutive samples of the MCMC chain. N $V_A$ <0.1 and N $h^2$<0.01 is the number of MCMC samples that estimated the additive genetic variance to be effectively zero and the narrow-sense heritability to be less than 0.01, respectively. r($GEBV_{default}$, $GEBV_{informed}$) is the correlation in GEBVs estimated from the default and informed prior models. Results are from models where all phenotyped and genotyped animals are used as the training population. Numbers on the second row of a cell indicate 95% credible intervals.*

| Variable | Weight | | Jaw length | | Foreleg | | Hindleg | | Metacarpal | | Horn length | | Coat colour | | Coat pattern | |
|---|---|---|---|---|---|---|---|---|---|---|---|---|---|---|---|---|
| Priors | Default | Informed | Default | Informed | Default | Informed | Default | Informed | Default | Informed | Default | Informed | Default | Informed | Default | Informed |
| Scaling parameter ($V_A$ $V_E$) | | 1.2, 2.5 | | 7.0, 6.8 | | 3.0, 9.0 | | 12.0, 14.0 | | 5.0, 5.4 | | 700, 1300 | | 0.09, 0.09 | | 0.01, 0.03 |
| $V_A$ | 1.67 | 1.90 | 10.00 | 10.98 | 7.57 | 7.87 | 16.74 | 18.01 | 8.10 | 8.37 | 1076 | 1352 | 0.061 | 0.062 | 0.006 | 0.007 |
| | 0.89-2.67 | 1.26-2.63 | 5.38-15.57 | 7.43-15.04 | 5.70-9.69 | 6.01-9.98 | 12.08-22.59 | 13.67-23.33 | 5.70-10.97 | 6.26-10.98 | 0-1835 | 815-1963 | 0.051-0.073 | 0.051-0.077 | 0.005-0.008 | 0.005-0.009 |
| $V_E$ | 3.03 | 2.92 | 8.37 | 7.99 | 7.45 | 7.39 | 17.06 | 16.63 | 4.73 | 4.76 | 1455 | 1250 | 0.066 | 0.066 | 0.028 | 0.028 |
| | 2.52-3.58 | 2.52-3.33 | 5.76-11.28 | 6.04-9.97 | 6.35-8.47 | 6.43-8.39 | 14.35-19.88 | 14.02-19.21 | 3.51-5.97 | 3.64-5.88 | 972-2784 | 966-1593 | 0.063-0.069 | 0.063-0.069 | 0.027-0.030 | 0.027-0.030 |
| $h^2$ | 0.36 | 0.39 | 0.54 | 0.58 | 0.50 | 0.52 | 0.50 | 0.52 | 0.63 | 0.64 | 0.43 | 0.52 | 0.48 | 0.48 | 0.18 | 0.19 |
| | 0.20-0.51 | 0.28-0.50 | 0.33-0.73 | 0.43-0.71 | 0.41-0.59 | 0.43-0.60 | 0.38-0.60 | 0.43-0.62 | 0.50-0.75 | 0.53-0.74 | 0.00-0.64 | 0.35-0.66 | 0.43-0.53 | 0.43-0.54 | 0.14-0.23 | 0.15-0.24 |
| N_SNPs | 3172 | 2540 | 4465 | 3297 | 1835 | 1136 | 2841 | 2729 | 4196 | 3133 | 1612 | 1198 | 535 | 502 | 430 | 343 |
| | 797-7570 | 428-5681 | 940-8402 | 769-6691 | 464-3877 | 237-3808 | 688-6595 | 494-5411 | 1059-7197 | 652-6225 | 259-4448 | 198-3420 | 130-967 | 153-930 | 57-1214 | 51-950 |
| N_SNPs_0.0001 | 2757 | 2297 | 4077 | 3017 | 1619 | 1138 | 2507 | 2509 | 3877 | 2829 | 1396 | 1036 | 488 | 454 | 379 | 288 |
| | 229-7453 | 129-5530 | 255-8325 | 350-6548 | 127-3730 | 32-3704 | 194-6487 | 124-5305 | 429-7101 | 115-6137 | 81-4375 | 37-3301 | 57-938 | 82-883 | 11-1174 | 7-908 |
| N_SNPs_0.001 | 376 | 168 | 360 | 216 | 149 | 114 | 291 | 153 | 292 | 253 | 163 | 95 | 24 | 21 | 33 | 29 |
| | 19-816 | 5-528 | 18-869 | 6-573 | 6-452 | 4-350 | 16-672 | 4-456 | 13-719 | 12-615 | 4-540 | 3-305 | 0-76 | 0-72 | 0-114 | 0-97 |
| N_SNPs_0.01 | 39 | 75 | 28 | 64 | 66 | 84 | 43 | 67 | 27 | 51 | 53 | 68 | 22 | 27 | 18 | 27 |
| | 2-92 | 30-121 | 1-76 | 24-109 | 33-102 | 48-121 | 12-86 | 30-107 | 6-60 | 21-93 | 11-123 | 34-104 | 14-35 | 16-41 | 8-34 | 13-46 |
| PGE_0.0001 | 0.28 | 0.23 | 0.42 | 0.31 | 0.16 | 0.12 | 0.25 | 0.25 | 0.39 | 0.29 | 0.15 | 0.11 | 0.05 | 0.05 | 0.04 | 0.03 |
| | 0.02-0.74 | 0.01-0.56 | 0.02-0.84 | 0.03-0.65 | 0.01-0.39 | 0.00-0.37 | 0.02-0.65 | 0.01-0.54 | 0.05-0.72 | 0.01-0.63 | 0.01-0.47 | 0.00-0.34 | 0.01-0.09 | 0.01-0.09 | 0.00-0.12 | 0.00-0.09 |
| PGE_0.001 | 0.38 | 0.17 | 0.36 | 0.22 | 0.15 | 0.11 | 0.29 | 0.15 | 0.29 | 0.25 | 0.15 | 0.09 | 0.02 | 0.02 | 0.03 | 0.03 |
| | 0.02-0.85 | 0.00-0.54 | 0.02-0.85 | 0.00-0.58 | 0.00-0.44 | 0.00-0.34 | 0.01-0.67 | 0.00-0.48 | 0.01-0.72 | 0.01-0.61 | 0.00-0.49 | 0.00-0.31 | 0.00-0.07 | 0.00-0.06 | 0.00-0.12 | 0.00-0.09 |
| PGE_0.01 | 0.34 | 0.60 | 0.22 | 0.48 | 0.68 | 0.78 | 0.45 | 0.60 | 0.31 | 0.46 | 0.70 | 0.80 | 0.92 | 0.92 | 0.92 | 0.91 |
| | 0.01-0.78 | 0.25-0.91 | 0.00-0.61 | 0.18-0.80 | 0.41-0.93 | 0.53-0.95 | 0.22-0.76 | 0.36-0.87 | 0.16-0.55 | 0.26-0.73 | 0.35-0.97 | 0.56-1.00 | 0.81-1.00 | 0.80-1.00 | 0.75-1.00 | 0.73-1.00 |
| Autocorr $V_A$ | 0.01 | 0.07 | 0.06 | 0.09 | 0.04 | -0.01 | -0.02 | 0.04 | 0.001 | 0.023 | 0.39 | 0.06 | 0.02 | 0.17 | 0.06 | 0.08 |
| N $V_A$ < 0.1 | 0 | 0 | 1 | 0 | 0 | 0 | 0 | 0 | 0 | 0 | 85 | 0 | 0 | 0 | 0 | 0 |
| N $h^2$ < 0.01 | 0 | 0 | 1 | 0 | 0 | 0 | 0 | 0 | 0 | 0 | 90 | 0 | 0 | 0 | 0 | 0 |
| r($GEBV_{default}$,$GEBV_{,informed}$) | | 0.9968 | | 0.9982 | | 0.9994 | | 0.9989 | | 0.9994 | | 0.9988 | | 0.9999 | | 0.9998 |

Figure S1: Chromosome partitioning plots of the eight traits. The y-axis shows the proportion of additive genetic variance explained by each chromosome

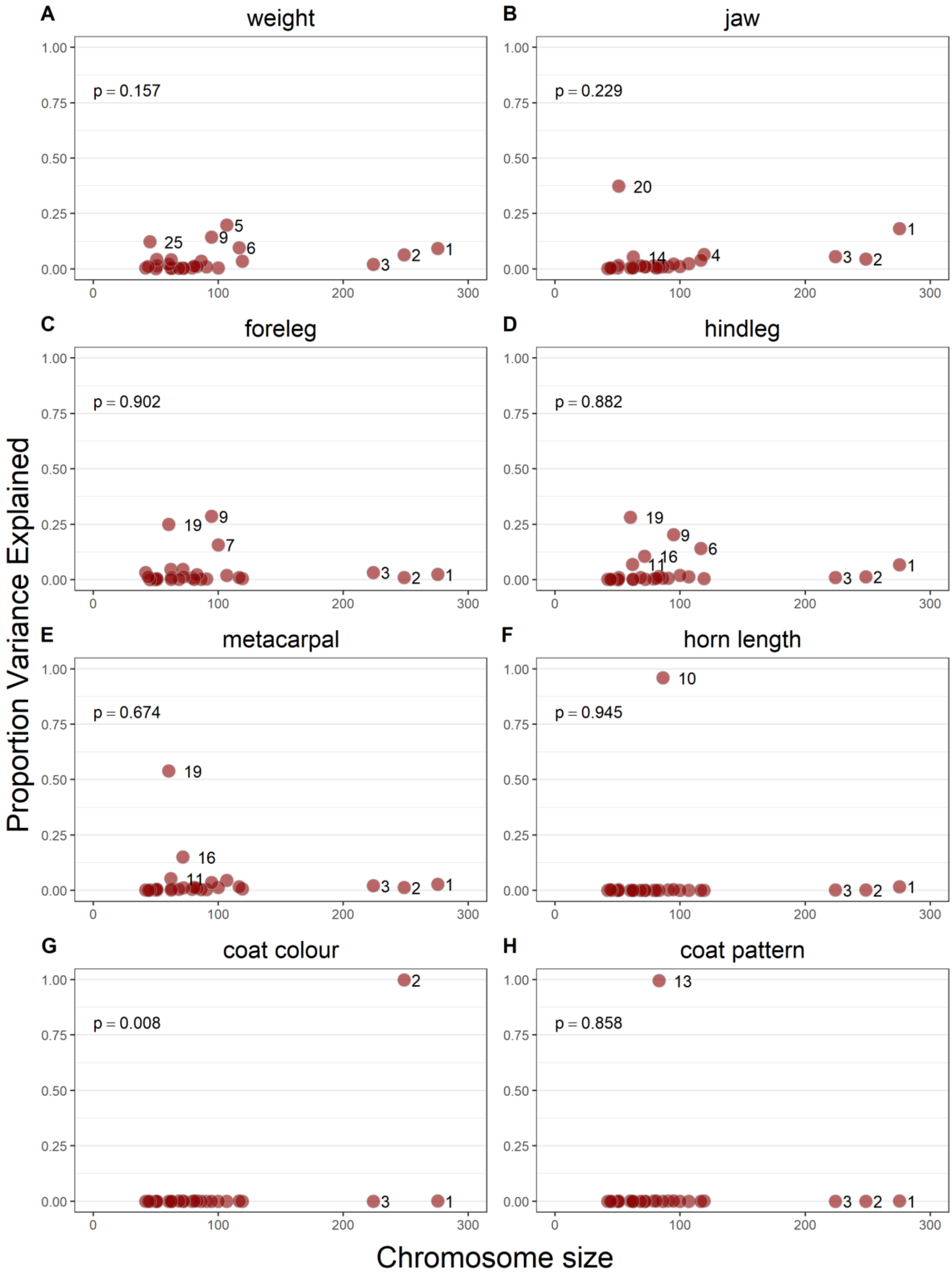

**Figure S2.** Manhattan plots of the eight traits. Y-axis shows the posterior inclusion probability of each SNP having an effect size >0 in the BayesR analysis. SNPs that have been identified in previous GWAS studies (referenced in Table 1 of manuscript) are shown as larger black symbols and labelled. SNPs with posterior inclusion probability >0.9 and that were not previously described in GWAS studies are labelled, but shown in smaller coloured symbols; e.g. for foreleg, SNP s48811.1

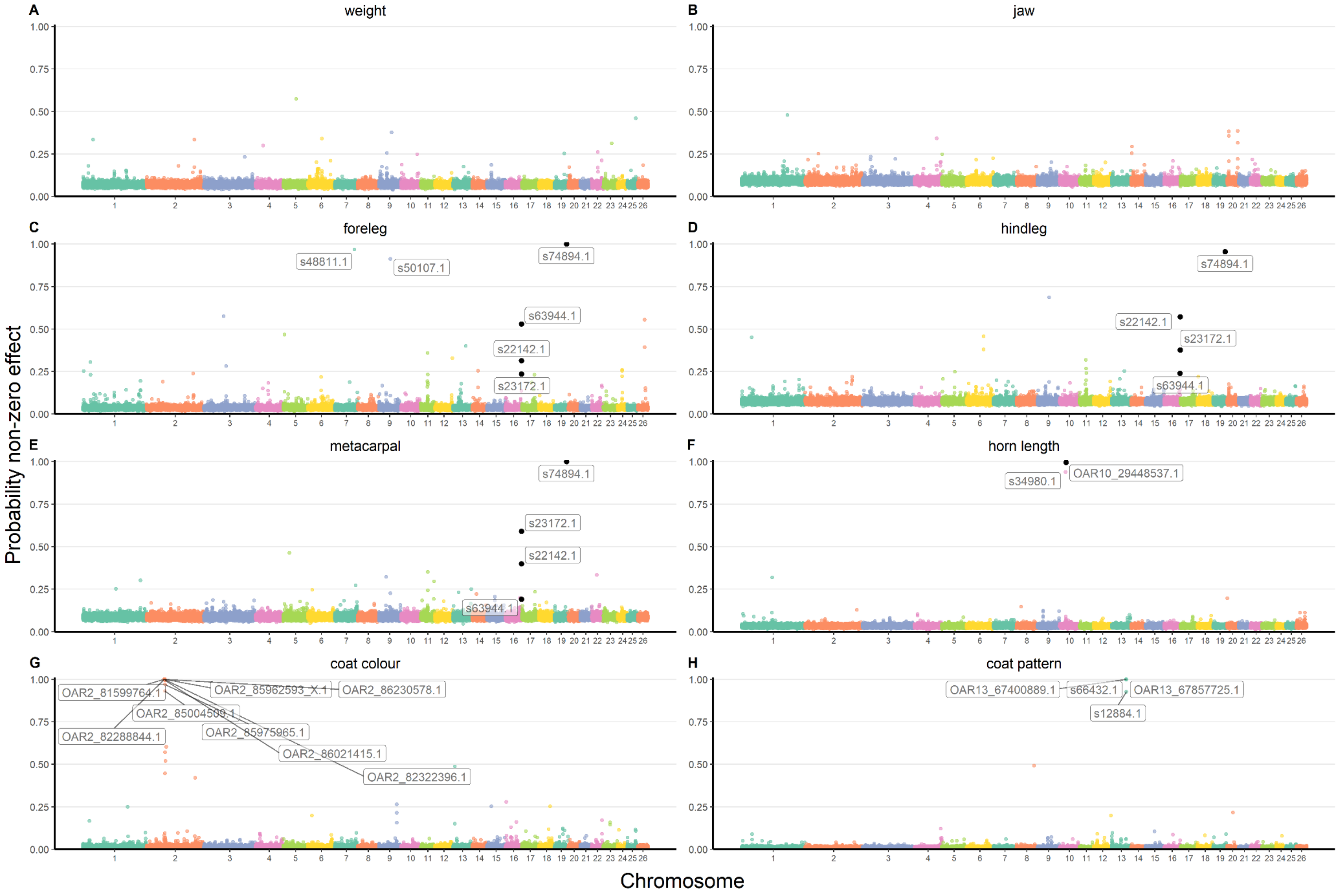

Figure S3. BayesR-derived genomic estimated breeding values for (A) coat colour and (B) coat pattern. Note the trimodal distributions, which reflects genotypes at the *TYRP1* (panel A) / *ASIP* (panel B) loci. At *TYRP1*, the dark allele is dominant to the light allele. At *ASIP* the wild type allele is dominant to the self pattern allele.

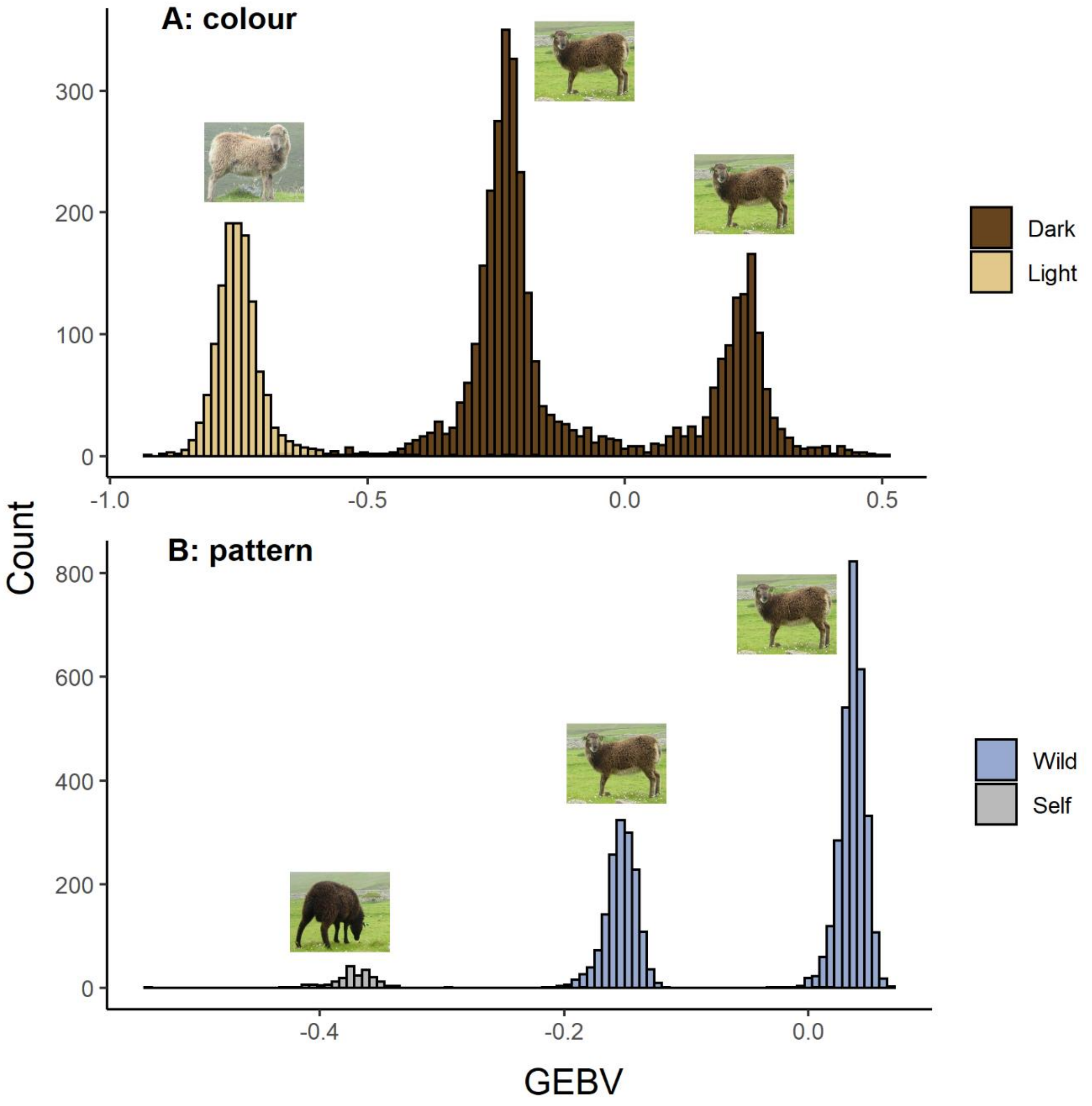
